# Non-driver somatic alteration burden confers good prognosis in non-small cell lung cancer

**DOI:** 10.1101/419424

**Authors:** Dennis Wang, Nhu-An Pham, Timothy M. Freeman, Vibha Raghavan, Roya Navab, Elena Pasko, Jonathan Chang, Chang-Qi Zhu, Dalam Ly, Jiefei Tong, Bradly G. Wouters, Melania Pintilie, Michael F. Moran, Frances A. Shepherd, Ming-Sound Tsao

**Affiliations:** Princess Margaret Cancer Centre, University Health Network, Toronto, Ontario, Canada; NIHR Sheffield Biomedical Research Centre, Sheffield, UK; Department of Laboratory Medicine and Pathobiology, University of Toronto, Toronto, Canada; Toronto General Research Institute, University Health Network, Toronto, Ontario, Canada; Program in Molecular Structure and Function, Hospital for Sick Children, Toronto, ON, Canada; Department of Medical Biophysics, University of Toronto, Toronto, Canada; Department of Medicine, University of Toronto, Toronto, Canada

**Keywords:** cancer genomics, prognostic biomarker, mutation burden, biomarker detection, patient stratification, cancer immunology, patient-derived xenograft

## Abstract

**Background:** Genomic profiling of patient tumors has linked somatic driver mutations to survival outcomes of non-small cell lung cancer (NSCLC) patients, especially for those receiving targeted therapies. However, it remains unclear whether specific non-driver mutations have any prognostic utility.

**Methods:** Whole exomes and transcriptomes were measured from NSCLC xenograft models of patients with diverse clinical outcomes. Penalised regression analysis was performed to identify a set of 865 genes associated with patient survival. The number of somatic copy number aberrations, point mutations and associated expression changes within the 865 genes were used to stratify independent NSCLC patient populations, filtered for chemotherapy naive and early-stage. In-depth genomic analysis and functional testing was conducted on the genomic alterations to understand their effect on improving survival.

**Results:** High burden of somatic alterations are associated with longer disease-free survival (HR=0.153, P=1.48×10-4) in NSCLC patients. When somatic alterations burden was integrated with gene expression, we were able to predict prognosis in three independent patient datasets. Patients with high alteration burden could be further stratified based on the presence of immunogenic mutations, revealing another subgroup of patients with even better prognosis (85% with >5 years survival), and associated with cytotoxic T-cell expression. In addition, 95% of these 865 genes lack documented activity relevant to cancer, but are in pathways regulating cell proliferation, motility and immune response were implicated.

**Conclusion:** Our results demonstrate that non-driver somatic alterations may influence the outcome of cancer patients by increasing beneficial immune response and inhibiting processes associated to tumorigenesis.

## BACKGROUND

The overall survival rate of lung cancer patients is poor at 10-20% ^1^; this is partly due to two thirds of the patients presenting in advanced stage disease at the time of diagnosis. However, even for one-third of the patients who potentially are curable by complete surgical resection, 50-80% will develop metastatic recurrence and die of the disease ^2^. Prognostic markers may be useful to identify patients with high risk of disease recurrence and death, and who may benefit from adjuvant chemotherapy ^3,4^. There have been extensive studies aimed at identifying prognostic markers for early stage NSCLC patients, including mutations in specific “driver” oncogenes and tumor suppressor genes, as well as gene expression signatures ^5–7^.

Whole-exome sequencing studies have suggested that the somatic alteration burden of tumors may also be prognostic of patient survival ^8–11^, possibly through the generation of immunogenic peptides known neoantigens ^12–15^. More recently, these biomarkers was able to predict patient response to immune checkpoint inhibitors ^16–18^. Despite these advances in establishing the prognostic value of mutational and neoantigenic burden, their value in early-stage NSCLC cases without the treatment of immunotherapies have yet to be explored. In this study, we analyse somatic aberrations found in patient-derived xenograft (PDX) tumors that are representative of patients ^19–22^, in order to evaluate the role and effect of mutation burden and neoantigen load on the prognosis of early stage completely resected NSCLC patients (Figure 1).

**Figure 1.**
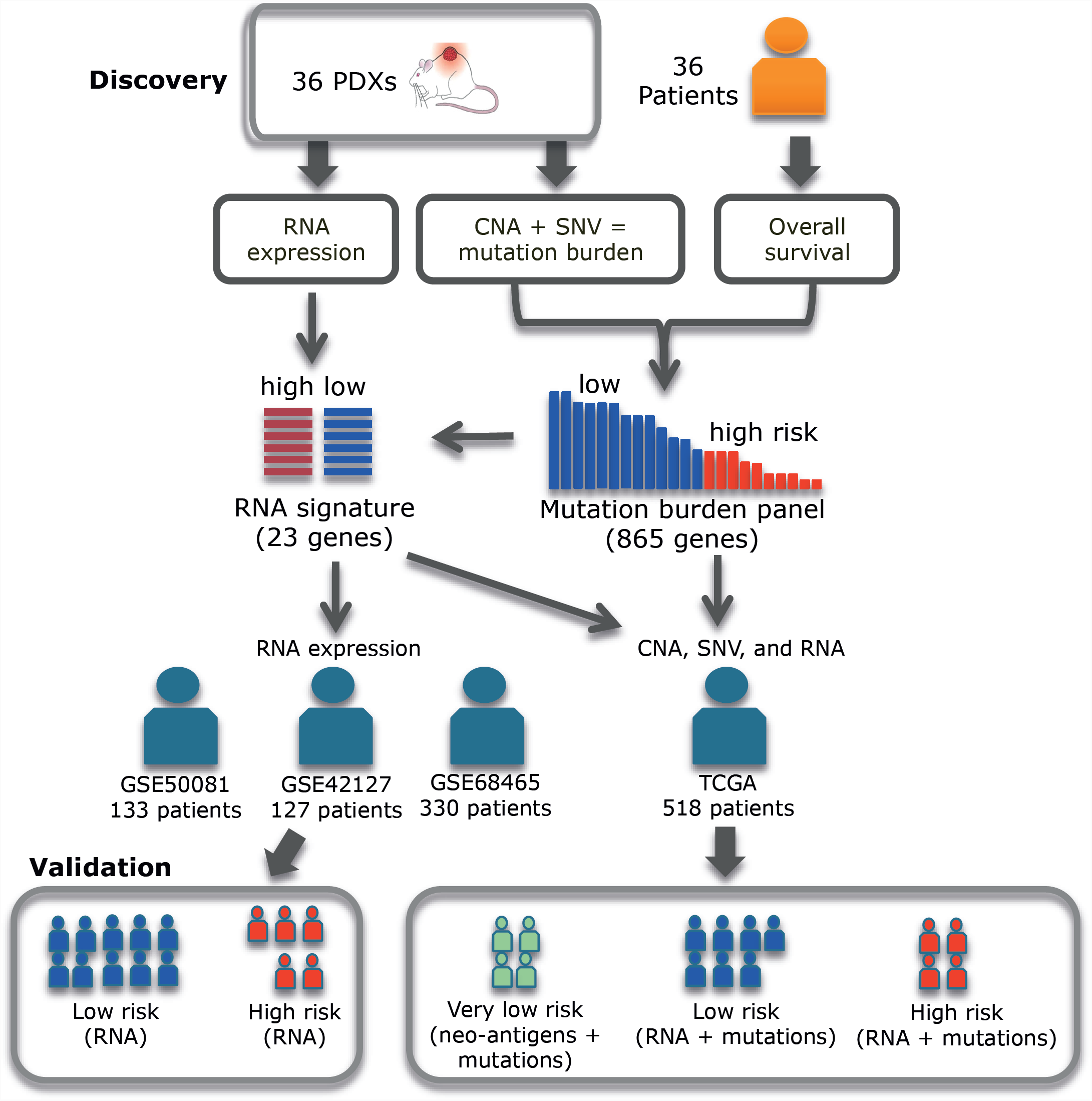
Translation of genomic information from PDXs to predict patient outcome. Somatic copy number (CNA), point mutations (SNV) and gene expression (RNA) profiles are measured in PDXs. A 23 gene expression signature and an 865 gene mutation panel was associated with survival of patients corresponding to the PDXs. The expression signature’s ability to predict OS was tested in three independent cohorts of NSCLC patients (GSE50081, GSE42127 and GSE68465) by stratifying patients into high and low risk of death. Survival prediction was also tested for 518 NSCLC stage I patients (TCGA) by integrating the expression signature, mutation panel and antigenicity of mutations measured in their tumors.

## METHODS

### Genomics data

Comprehensive whole-exome and transcriptome profiles from 36 PDXs, their matched primary tumors and normal tissue were obtained ^19–22^. TCGA mutation, copy number, RNA-seq, and clinical data were downloaded for LUAD ^23^ and LUSC ^24^ patients on cBioPortal, using the cgdsr R package ^25^. Preprocessing was performed consistently for both datasets (Supplementary Methods).

### Stratification by mutation burden

To identify genes associated with patient survival from PDX molecular profiles, we used ElasticNet penalised regression to fit gene-level copy number aberrations and mutations (Supplementary Methods). The number of alterations among 865 genes (NAG) was used as a risk score for each TCGA patient in the validation set. The risk scores were dichotomized at the first quartile to assign patients to one of the prognostic arms (high vs. low number of alterations). Proportion of disease free survival (DFS) and overall survival (OS) was calculated using the Kaplan-Meier method and the difference between curves was tested using the log-rank test. OS is the time between the date of diagnosis to the date of death or last follow-up, and DFS is the date of diagnosis to the date of death or relapse or last follow-up.

### Stratification by gene expression

A gene expression signature associated with NAG is composed of 79 genes differentially expressed in high vs low NAG PDXs. The UT Lung Spore dataset (GSE42127)^26^ was then used to further select genes associated with patient survival (Supplementary Methods). Validation of NAG expression signature was performed on the Director’s Challenge Consortium for Molecular Classification of Lung Adenocarcinoma data set with a total of 442 samples (GSE68465), and the UHN181 cohort of 181 NSCLC patients (GSE50081). The risk scores from the expression signature with NAG classification to predict survival of TCGA patients. A patient classified as high risk from the expression signature and low NAG would be predicted to be poor prognosis.

### Stratification by immunogenicity

Non-synonymous mutations causing amino acid changes were identified among the 865-gene panel for TCGA patients. We assessed the MHC-I binding affinity of various 8-11mer amino acid sequences containing the mutation using NetMHCpan v 3.0 ^27^ (Supplementary Methods). We defined peptides as antigenic if they had an HLA-A binding affinity below 500nM with both HLA-A alleles. In addition, antigenic peptides’ genes had to be expressed above the median expression level across all genes and patients in order for the peptides to be considered immunogenic, and the patient HLA-A gene needed to be expressed above the median HLA-A expression level across all patients. Patients were stratified based upon whether or not they had any immunogenic peptides as outlined above, either by itself, or in combination with the NAG score.

### Pathway enrichment

Gene set enrichment was performed on the 865 genes using the DAVID and WebGestalt^28^. Pathway memberships of genes were extracted from GeneCards, KEGG, and PathwayCommons ^29^. Cytoscape (v3.2.0) was used to customize and visualize the network. Nodes on the edges of the network that did not connect to any of the main cancer pathways were removed (i.e. CLTC and EPN1). The genes were arranged by cellular location into either the ECM and plasma membrane or the cytoplasm.

### In vitro functional assays

Cell proliferation assays were performed on A549, H1573 and HCC827 cells as described previously ^30^. In brief, 5,000 cells were seeded per well of E-plate. Impedance was measured every 15 minutes for 80 hours. Growth curves were constructed using the xCELLigence platform (ACEA Biosciences, San Diego, CA). Tumor cell invasion was assessed *in vitro* by the reconstituted basement membrane (Matrigel) invasion assay ^31^, which was performed using 8-μm polycarbonate filters coated with reconstituted basement membrane (Matrigel; BD Bio-sciences). Motility of cells were measured after seeding in a plastic-bottomed 24-well dish and incubated for 12 hours. Images were acquired every 20 minutes for approximately 24 hours using a 10x phase objective. Further conditions are described in the Supplementary Methods.

### Statistical Analysis

All reported hazard ratios (HR) scores and p-values use Kaplan-Meier method unless reported as multivariate analyses. A multivariate Cox proportional hazards model was used to fit survival times to the number of altered gene while adjusting for other clinical factors including age, stage, and smoking status. All reported survival differences within the Cox model were tested using the Wald test, and follow-up of patients were measured up to 5 years. All other statistical tests used to measure difference in quantities are stated and two-sided.

## RESULTS

### Somatic Alteration Burden in Xenografts is Correlated with Patient Survival

Tumors from 36 NSCLC patients were engrafted and successfully passaged in immune-deficient mice to form PDXs ^19,21^. The number of alterations (CNA or mutation) per PDX varied from 164 to 765 (Figure 2A). By comparing the DFS and OS of patients of PDXs with ≤ 343 alterations (first quartile) to patients of PDXs with >343 alterations, group with the higher number of alterations had significantly better OS (HR= 0.366, 95% CI=0.140-0.952, P=0.040) and DFS (HR=0.294, 95% CI=0.103-0.840, P=0.022). The comparison of CNAs alone with DFS showed weaker and non-significant association (HR=0.546, 95% CI=0.185-1.61, P=0.272); this also applies to OS (HR=0.532, 95% CI=0.197-1.43, P=0.202). Similarly, the number of somatic mutations alone was not significantly associated with DFS (HR=0.568, 95% CI=0.179-1.79, P=0.335) or OS (HR=1.41, 95% CI=0.407-4.76, P=0.592). When we examined gene expression and protein phosphorylation for the same phenomenon, we observed only a moderate association (*r*=0.32, P=0.087) between the number of proteins that were tyrosine phosphorylated and the number of genes altered across PDXs, but the number of phosphorylated proteins remained positively correlated with DFS (Supplementary Figure 1).

**Figure 2.**
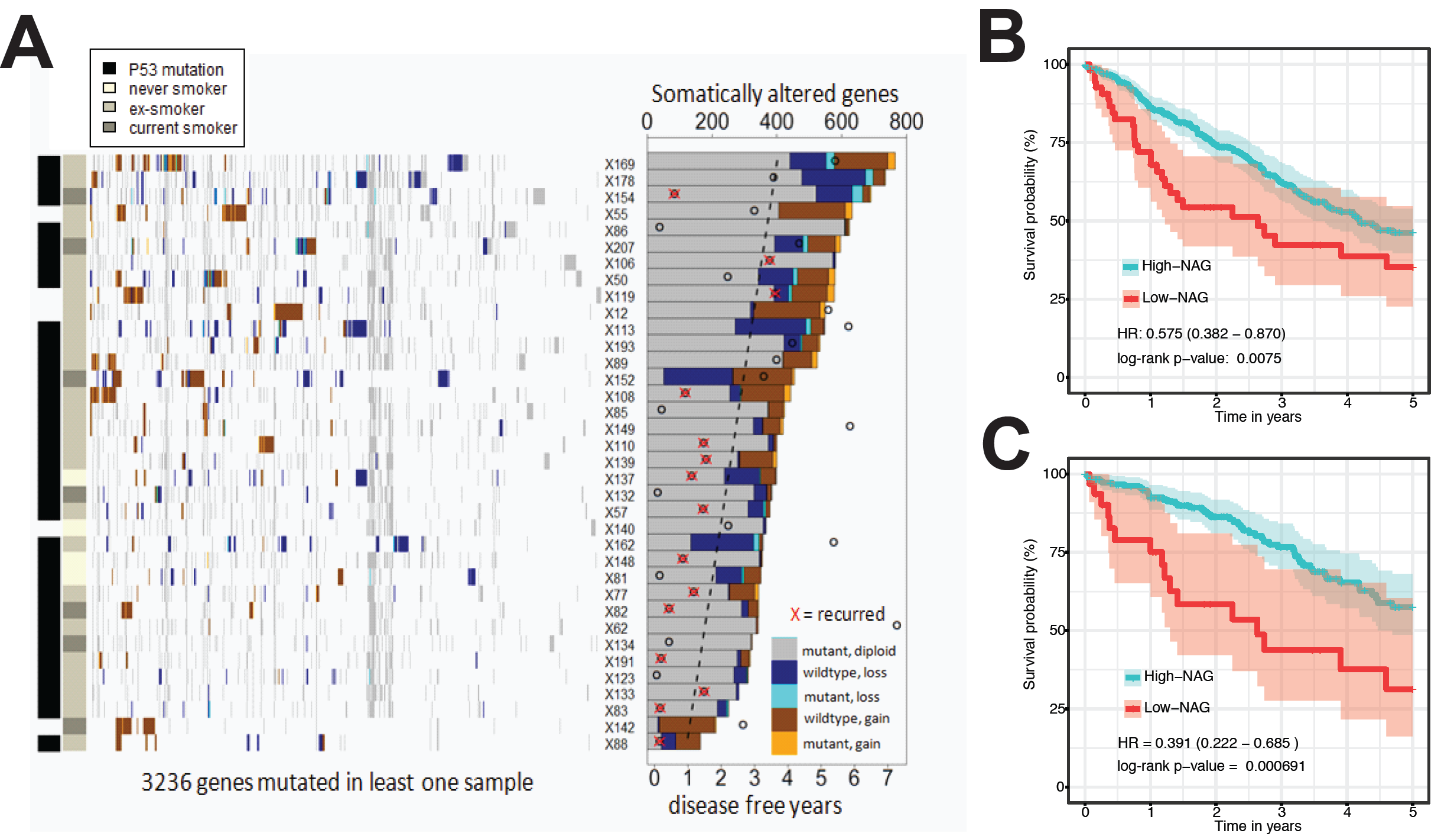
Association between somatic alterations found in PDXs and OS of patients. (A) A heatmap of somatic mutated and copy number altered genes across PDX models. The total NAG in each PDX is described by the bar graph. Circles on the bar graph show the disease-free years for each PDX’s matched patient, and a dashed line shows best fit running through them. Genes are ordered according to hierarchical clustering across samples. Kaplan-Meier curves show significant difference (log-rank test) in OS between high and low NAG in (B) all TCGA NSCLC patients and (C) only in stage I patients.

A signature for OS was generated based on genomic alterations across the 3,236 genes using sparse regression. Genes that were altered in only one PDX were removed, and then the status of each gene was dichotomized as altered or not altered. After fitting the alteration profiles to patient survival using penalized regression, 865 genes were identified with non-zero coefficients that contributed the most to overall risk of death (Supplementary Table 1). Only 47 of these genes (5.4%) had documented activity relevant to cancer according to the Cancer Gene Census ^32^, these included DNA repair genes TP53 and BRCA1. Based on the alteration status of these 865 genes, the PDXs were divided into two groups using a cut-off at the first quartile (88 genes) for NAG. Patients with a high NAG in their PDXs had better survival (HR=0.153, 95% CI=0.051-0.459, P=1.48×10^−4^) than patients whose PDXs had a low NAG (Supplementary Figure 2).

### Alteration burden predictive of survival for early stage NSCLC

Since somatic alterations affect OS, we hypothesized that there might be differentially expressed genes between the high and low alterations groups that also might be prognostic. We identified 79 genes with differentially expressed mRNA (> 2 absolute fold change, P < 0.05) between the two groups of PDXs based on the NAG (Supplementary Table 2). Using the mRNA expression data of the 79 genes from 127 NSCLC patients (GSE42127)^17^, expression scores associated with survival were calculated. Among the 79 genes, the expression of only 23 genes had non-zero coefficients in the regression with OS in this patient cohort. A NAG expression signature was developed based on 23 out of the 79 genes that had the greatest effect on survival. In the training set of 127 patients, this NAG expression signature was significantly prognostic (Supplementary Figure 2; HR 0.23, 95% CI 0.125-0.422, P=2.3×10^−7^). This was validated in two independent early stage, untreated patient cohorts with microarray data (GSE50081: HR=0.562, 95% CI=0.332-0.952, P=0.029; GSE68465: HR=0.518, 95% CI=0.338-0.794, P=2.2×10^−3^; Supplementary Figure 2).

To test the alteration burden together with the expression signature as a prognostic classifier for NSCLC, we attempted to predict overall survival of 221 TCGA LUAD and 173 LUSC cases. We counted the NAG among the 865 genes in each patient tumor, then classified the patients using the same cut-off (88 altered genes) that was used for classifying high and low altered-gene PDX tumors, and the expression risk score from RNAseq data from patient tumors. All patients with tumors containing ≥88 altered genes or a low expression risk score had significantly better OS compared to those with <88 altered genes and a high expression risk score (Figure 2B; HR=0.575, 95% CI= 0.382 −0.870, P=0.0075); this also was true for stage I patients (Figure 2C; HR=0.391, 95% CI= 0.222 - 0.685, P=6.9×10^−4^). Multivariate survival analyses of stage I patients adjusted for age, sex, smoking history, and histology confirmed that the number of alterations was an independent predictor of good OS (HR=0.407, 95% CI= 0.211 - 0.787, P=7.7×10^−3^, Table 1).

**Table 1:** Multivariate survival model of NAG score with clinicopathological factors of NSCLC Stage I patients.

|  | HR | 95% CI | Wald test P-value |
| --- | --- | --- | --- |
| Age( >65 vs. <=65) | 1.04 | 0.62-1.73 | 0.89 |
| Sex (F vs. M) | 0.85 | 0.51-1.43 | 0.54 |
| Tobacco (smoker vs never) | 1.75 | 0.74-4.12 | 0.2 |
| Histology (Adeno vs. Squamous) | 0.96 | 0.52-1.79 | 0.91 |
| Overall NAG score (low vs high burden) | 2.46 | 1.27-4.75 | 0.0077* |

*NAG* number of altered genes, *HR* hazard ratio, *CI* confidence interval

### Mutated immunogenic peptides correlate with best survival and cytotoxic T-cell signatures

Given that alteration burden is associated with good survival outcome, we hypothesized that some of the mutations in the 865 genes may induce an immune response towards the tumor through the expression of immunogenic peptides or neoantigens. This was tested by examining whether somatic SNV within the 865 genes found in TCGA NSCLC patients could encode for tumor specific antigens presented by MHC Class I molecules; 86 patients had at least one expressed mutated peptide with high MHC I binding affinity against both *HLA-A* alleles and this immunogenic group had significantly better OS (Supplementary <u>Figure 3A</u>; HR=0.536, CI=0.341 - 0.8419, P=0.0068). Among stage I patients, 47 were classified with immunogenic tumors and they also had significantly better OS (Figure 3A; HR=0.266, CI=0.1068 - 0.6619, P=0.0044, even after adjusting for other clinical factors (Table 2). In comparison, patients stratified using all neoantigens identified, beyond those in the 865 panel of genes, had no significant difference (P < 0.01) in overall survival (Supplementary <u>Table 3</u>). By combining this immunogenicity classifier with the alteration burden classifier based on NAG, we were able to identify good and poor prognosis patients. Stage I patients with both immunogenic and high NAG tumors had significantly better survival than patients with both low NAG and no neoantigens (Figure 3B; HR=0.0807, CI=0.02315 - 0.2813, P=7.8e-05). The RNA expression profiles from these two groups of patients were further analysed for immune components. Immune cell population estimates from TIMER ^33^ revealed moderately higher proportions of B-cells, dendritic cells, and CD8 T-cells in high NAG patient tumors with immunogenic peptides compared to without (Figure 3C). Focusing specifically on markers of cytotoxic T-cells in Figure 3D, we also found in high NAG tumors with immunogenic peptides significantly higher (P < 0.05) expression of CD8A, granzyme B (GZMB) and perforin (PRF1). There was also significantly higher expression of cytokines CCL5, CXCL9, CXCL10, and IL16 that are associated with cytotoxic T-cells ^34^. Patients with no immunogenic peptides were also stratified into high and low NAG groups, but there were no concordant difference across multiple immune cell populations or cytotoxic T-cell markers (Supplementary Figure 3B,C).

**Table 2:** Multivariate survival model of immunogenicity factor with clinicopathological factors of NSCLC stage I patients.

|  | HR | 95% CI | Wald test P-value |
| --- | --- | --- | --- |
| Age of diagnosis ($\leq 65$ yrs vs $> 65$ yrs ) | 0.929 | 0.545 - 1.583 | 0.7865 |
| Gender (male vs female) | 0.8705 | 0.524 - 1.447 | 0.5929 |
| Smoking history (last 15 yrs; yes vs no) | 0.7514 | 0.434 - 1.302 | 0.3085 |
| Histology (adeno. vs squamous cell) | 0.6767 | 0.392 - 1.169 | 0.1615 |
| Immunogenicity ($\geq 1$ neoantigen vs 0 neoantigens) | 0.2961 | 0.119 - 0.740 | 0.00919 * |

*HR* hazard ratio, *CI* confidence interval

**Figure 3.**
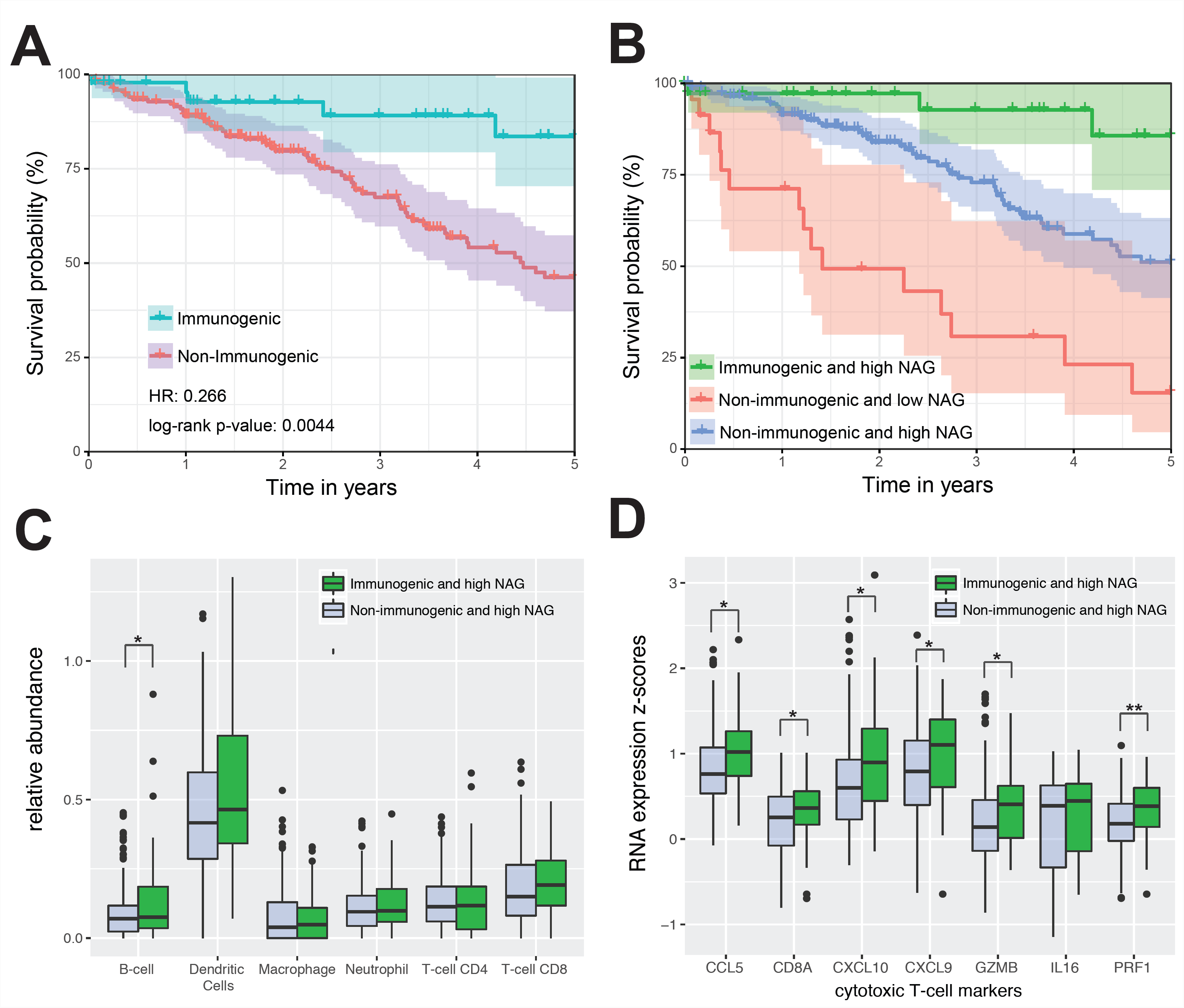
TCGA NSCLC patients stratified by immunogenic neoantigens. (A) Stage I patients (47 with neoantigens, 171 without neoantigens), are grouped into those with immunogenic neoantigens and those with none. Hazard ratios and log-rank p-values compare immunogenic patient tumors to non-immunogenic patient tumors. (B) Overall survival difference of stage I patients classified based on presence of immunogenic neoantigens and the number of altered genes. Immunogenic and high-NAG patient tumors (green, 37 patients) vs non-immunogenic and low-NAG patient tumors (red, 23 patients) show HR= 0.0807 P=7.8e-05. Immunogenic and high-NAG tumors (green) vs non-immunogenic and high-NAG patient tumors (blue, 147 patients) show HR= 0.229 P=0.013. Non-immunogenic and high-NAG tumors (blue) vs non-immunogenic and low-NAG patient tumors (red) show HR= 0.309 P=7.9e-05. High NAG patients with and without neoantigens were contrasted based on (C) the relative abundance of immune cell types estimated by the TIMER algorithm, and (D) the RNA expression of cytotoxic T-cell markers. Significant differences for each component are marked (* t-test p-value < 0.05; ** < 0.01).

### Prognostic Alterations Enriched in Extra-Cellular Signaling

In order to investigate the mechanism by which alteration burden may be impeding cancer progression, pathway enrichment analyses were performed on the 865 set of genes associated with alteration burden. The most enriched pathways involved BARD1 signaling, Janus kinase activity, integrin signaling and extra-cellular matrix interactions (Figure 4; hyper-geometric test P<0.01). Within these pathways, a number of genes were altered more frequently across TCGA cases in the group of low-risk patients with better survival (Supplementary Figure 4). Inactivating alterations, which deactivate Wnt signaling, were most frequent in *CTNNB*1 (23.1%) and *WNT5A* (22.8%), along with copy gains in tumor suppressors *NHERF1* (16.4%) and *NF2* (13.1%). High frequency of copy gains occurring in *ITGA10* (20.9%), *COL20A1* (17.9%) and *COL8A1* (26.5%) suggested that cell adhesion to the extracellular matrix may be affected.

**Figure 4.**
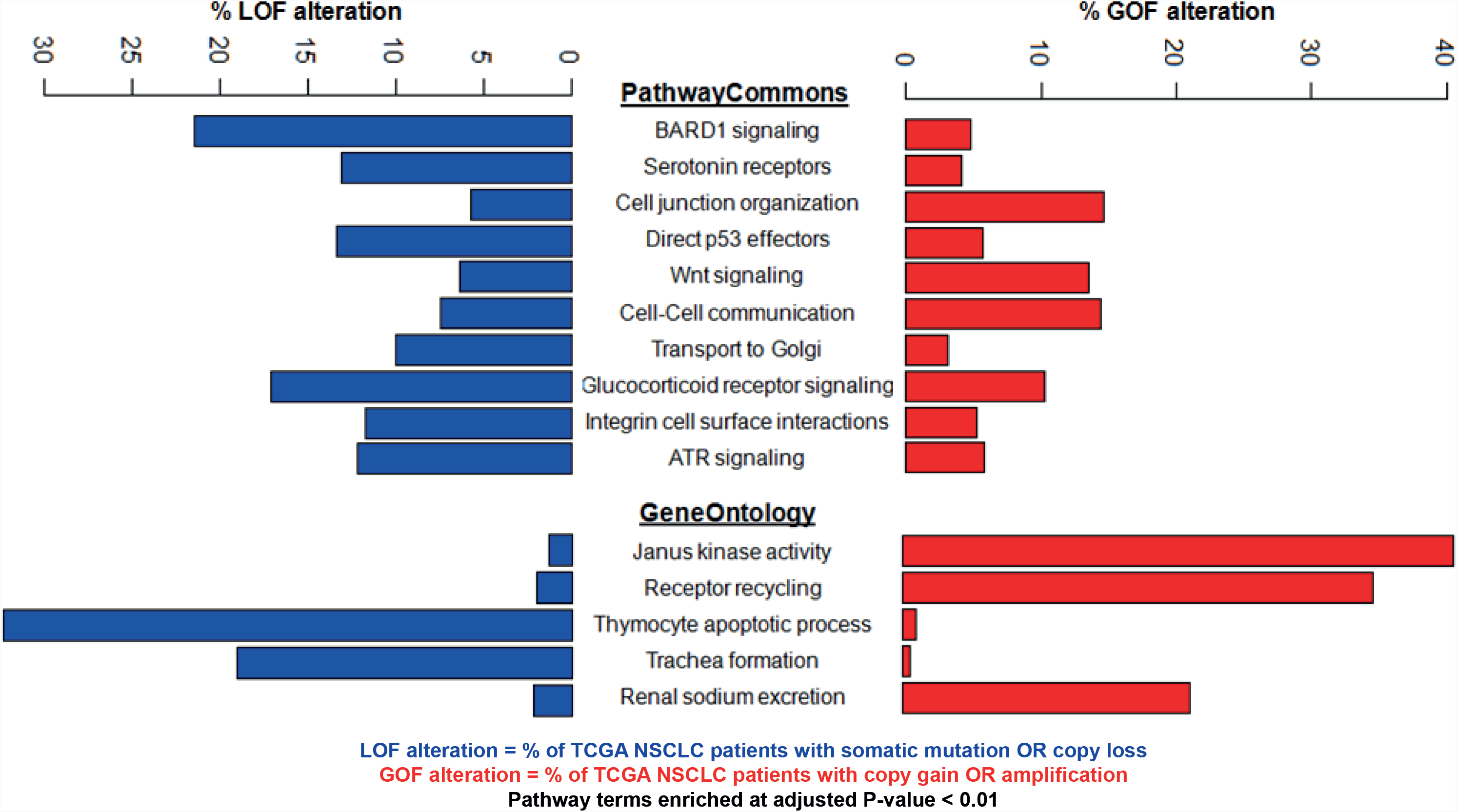
Enriched pathways and biological processes for genes in NAG. Barplots show proportion of genes with somatic alterations that are classified as gain of function (GOF) or loss of function (LOF).

To evaluate the effect of genomic alterations on the proteome, we profiled the phospho-protein profiles of high NAG tumors and identified known cancer associated genes with elevated levels of tyrosine phosphorylation (Supplementary Table 4). We also evaluated the functional impact of mutation burden by selecting three NSCLC cell lines with different risk scores as determined by their NAG and their NAG expression scores (Figure 5A). The cell line (HCC827) with the highest estimated overall risk score had the highest rate of proliferation, cell motility and invasion compared to the other two cell lines, H1573 and A549 (Figure 5B-D).

**Figure 5.**
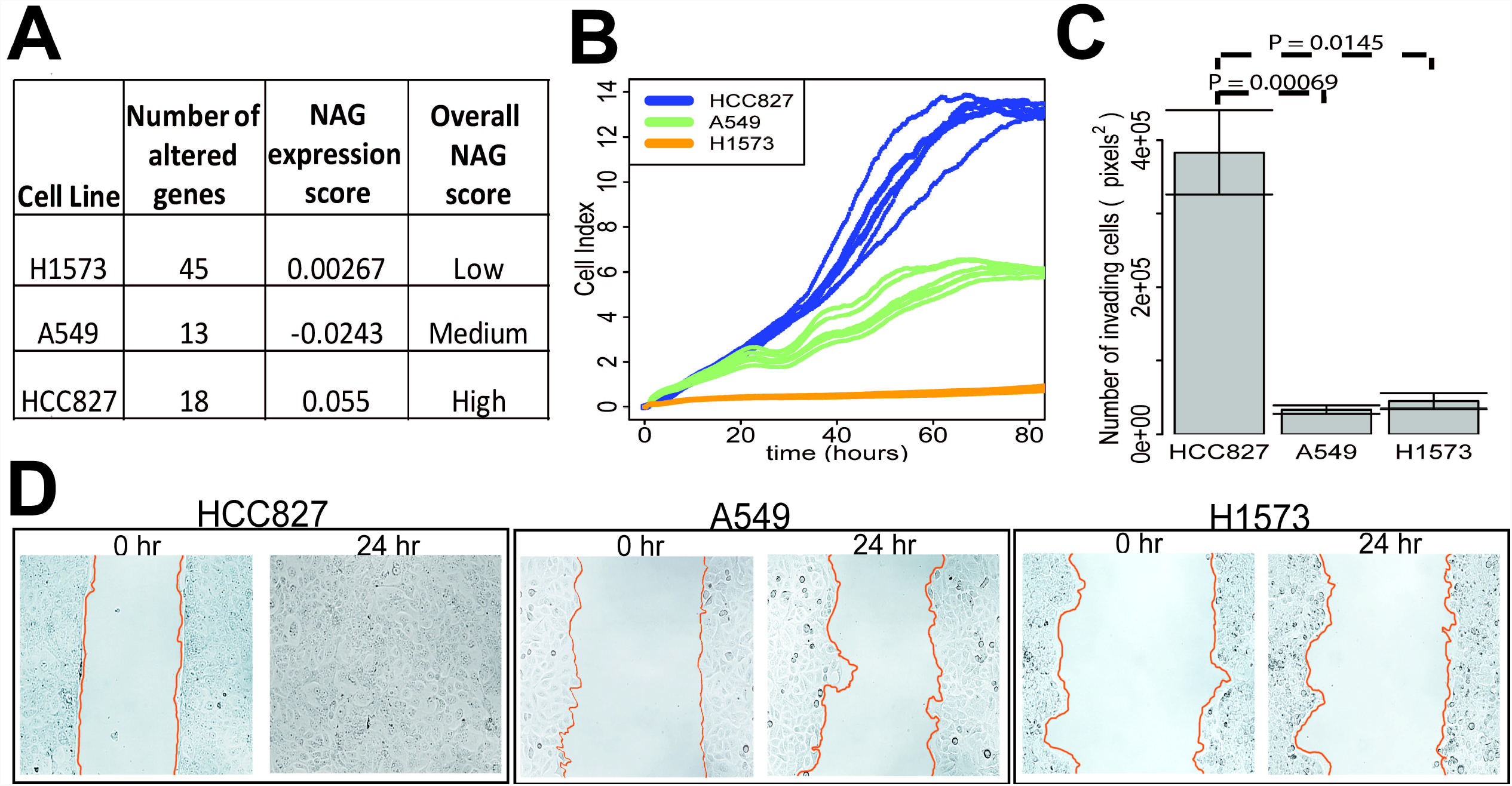
Functional impact of altered and differentially expressed genes on cancer cells. (A) Three lung cancer cell lines with differing risk scores according to the NAG and NAG associated expression signature were compared to each other in functional assays. (B) Proliferation of the three cultured cell lines was estimated by electrical impedance (cell index) over time. (C) Invasion of cells across transwell membranes coated with Matrigel was counted. (D) Cell motility was measured by a time-lapse wound-healing assay. The rates of wound area travelled by cells were 15.42µm/hr for HC827, 2.64µm/hr for A549, and 1.98µm/hr for H1573.

## DISCUSSION

We have shown that an estimate of somatic alteration burden among 865 genes derived from PDX tumor models is predictive of prognosis for early stage NSCLC cases. This integrative burden estimate was correlated with the survival of PDX patients and predicted the survival of early stage TCGA patients and untreated patients in two other independent cohorts. In particular, patients with low alteration burden had significantly poorer OS than those with high alteration burden. Of the patients with high alteration burden, there was a subset with immunogenic point mutations that had significantly better survival than other stage I patients. Pathway analysis and functional investigation revealed that 95% of the 865 genes were not in cancer drivers, but they are associated with processes relating to immune activation, extracellular matrix interactions, and cell motility.

Our findings showed that poor OS in early stage NSCLC can be predicted by low somatic alteration burden in a select set of 865 genes. Previous studies examining mutation or genetic alteration burden in lung cancers have found the opposite effect of high burden being associated with poor survival ^35,36^, however, these relied on profiling whole exomes and targeted cancer driver events. In contrast, we performed targeted analyses of exons in only 865 genes, of which very few were previously associated with cancer progression. Smaller targeted gene panels have recently been shown to be effective at measuring overall tumor mutation burden and have been applied as a companion diagnostic ^37^. The mechanism by which alteration burden affects cancer progression and patient survival remain unclear, but it has been hypothesized that excessive genomic instability can lead to deleterious mutations that impedes tumor growth ^38^. Mutations in DNA damage response genes, such as *TP53* and *BRCA1*, which were among the 865 gene panel are markers of elevated mutation rate ^39,40^. Greater DNA damage could result in deleterious mutations or copy number losses in many of the other 865 genes that are associated with cell adhesion, motility and integrin signaling. Integrin signaling is a major pathway contributing to cancer cell survival and has been a target for antagonists to inhibit tumor growth ^41^. The lack of somatic alterations in integrin pathway genes may also be a marker for aggressive tumors in patients with poor survival outcomes. This could explain why cell lines, such as HCC827, with low alteration burden was able to proliferate and invade rapidly.

Somatic mutation burden has been correlated with neoantigen load and T-cell infiltration in NSCLC and melanoma ^10,42,43^, yet no association has been found in early-stage cancers where there is a lower mutation rate and which are predominantly untreated with therapeutics. Our assessment of neoantigens among the 865 genes shows that even without immunotherapy, the presence of neoantigens and high mutation burden is associated with cytotoxic T-cells in early-stage NSCLC. Previously mutation burden and neoantigen load have been used to identify patients with good prognosis following immunotherapy treatment, such as immune checkpoint inhibitors, but there has been debate about whether both markers are needed given that they select for a similar subgroup of patients ^37,42,44^. We showed that a combination of low alteration burden and no neoantigens had far worse OS (median survival of 2 years).

By translating findings from PDX models, we were able to identify two prognostic markers of survival for stage I NSCLC patients, their somatic alteration burden and the presence of immunogenic peptides. We found utility in using both alteration burden and neoantigen load for informing prognosis. This work also encourages further research to be done to understand the effect of mutations in non-cancer driver genes and the impact of immune activity in early cancer development.

## ACKNOWLEDGEMENTS

D.W., N.-A.P., F.A.S., B.G.W., and M.S.T conceived the study and devised the experiments. D.W., N.-A.P., T.M.F., J.C., C.-Q.Z, D.L., M.P. and V.R are involved in data processing, analysis and interpretation. R.N. and E.P. performed functional testing and analysis on cell lines. D.W. and M.S.T. wrote the first draft of the paper. All other co-authors contributed to the writing and have approved the final manuscript. We wish to thank Jenna Sykes and Ni Liu for assistance.

